# Alkaloid and acetogenin-rich fraction from *Annona crassiflora* fruit peel inhibits proliferation and migration of human liver cancer HepG2 cells

**DOI:** 10.1101/2021.04.07.438799

**Authors:** Allisson B. Justino, Rodrigo M. Florentino, Andressa França, Antonio C. M. L. Filho, Rodrigo R. Franco, André L. Saraiva, Matheus C. Fonseca, Maria F. Leite, Foued S. Espindola

## Abstract

Plant species from Annonaceae are commonly used in traditional medicine to treat various cancer types. This study aimed to investigate the antiproliferative potential of an alkaloid and acetogenin-rich fraction from the fruit peel of *Annona crassiflora* in HepG2 cells. A liquid-liquid fractionation was carried out on the ethanol extract of *A. crassiflora* fruit peel in order to obtain an alkaloid and acetogenin-rich fraction (AF-Ac). Cytotoxicity, proliferation and migration were evaluated in the HepG2 cells, as well as the proliferating cell nuclear antigen (PCNA), vinculin and epidermal growth factor receptor (EGFR) expression. In addition, intracellular Ca^2+^ was determined using Fluo4-AM and fluorescence microscopy. First, 9 aporphine alkaloids and 4 acetogenins that had not yet been identified in the fruit peel of *A. crassiflora* were found in AF-Ac. The treatment with 50 µg/mL AF-Ac reduced HepG2 cell viability, proliferation and migration (*p* < 0.001), which is in accordance with the reduced expression of PCNA and EGFR levels (*p* < 0.05). Furthermore, AF-Ac increased intracellular Ca^2+^ in the HepG2 cells, mobilizing intracellular calcium stores, which might be involved in the anti-migration and anti-proliferation capacities of AF-Ac. Our results support the growth-inhibitory potential of AF-Ac on HepG2 cells and suggest that this effect is triggered, at least in part, by PCNA and EGFR modulation and mobilization of intracellular Ca^2+^. This study showed biological activities not yet described for *A. crassiflora* fruit peel, which provide new possibilities for further *in vivo* studies to assess the antitumoral potential of *A. crassiflora*, especially its fruit peel.

## Introduction

Cancer represents one of the main challenges for medicine being one of the most critical problems of public health in the world. Hepatocellular carcinoma (HCC) is the seventh most frequently occurring cancer and the fourth most common cause of cancer mortality, with over half a million new cases diagnosed annually worldwide [1]. Hepatitis B and C virus and excessive alcohol consumption are important risk factors for HCC [2]. In addition to its high incidence, this tumor is usually diagnosed at advanced stages, which hampers effective treatment. Thus, the search of new agents capable of controlling the development of hepatocellular tumor is important to reduce the mortality caused by this disease.

Thus, over the past decade, numerous studies have shown that compounds derived from plants are potentially interesting for therapeutic interventions in various cancer types due to their great diversity of chemical structures, and better drug-like properties compared to the synthetic compounds [3–5]. Examples include plant-derived alkaloids, specifically aporphine alkaloids, which had previously demonstrated antitumor effects in different cancer cell models [5–8]. Acetogenins, a class of polyketide compounds found in plants of the Annonaceae family, have also been reported to possess apoptosis-inducing effects [9]. *Annona crassiflora* Mart., an Annonaceae species common to the Brazilian Savanna, where it is known as araticum, might be a potential source of acetogenins and aporphine alkaloids [10–12]. Different parts of this species such as bark, leaf, fruit and seed have been widely used in folk medicine for the treatment of inflammation, microbial infections, malaria, veneral diseases, snakebites, diarrhea, and as cancer chemopreventive agents [13–15].

Recently, methanolic extracts of leaves and seeds of *A. crassiflora* have shown *in vitro* antiproliferative properties in leukemia, glioblastoma, lung and ovarian cancer cell lines [16]. Furthermore, a study done by Silva, Alves (17) showed that a hexane fraction from the crude extract of *A. crassiflora* leaf had cytotoxic effect on cervical cancer cells by acting through DNA damage, apoptosis via intrinsic pathway and mitochondrial membrane depolarization [17]. However, scientific reports demonstrating antitumoral activities of the fruit peel of this species are still limited.

Previously, a pre-purification of the ethanol extract of *A. crassiflora* fruit peel was conducted, resulting in an alkaloid (CH_2_Cl_2_ fraction)-enriched fraction [10]. From the CH_2_Cl_2_ fraction, stephalagine, an aporphine alkaloid, was isolated and characterized [11]. It is worth mentioning here that the only biological activities described for this alkaloid are its antinociceptive feature [10] and potential inhibitory effect against pancreatic lipase [11]. In this context, in this study, we first identified the main alkaloids and acetogenins present in the alkaloid and acetogenin-rich fraction from *A. crassiflora* fruit peel, named here as AF-Ac. Then, we evaluated the antiproliferative potential of AF-Ac in HepG2 cells, exploring the possible involvement of the proliferating cell nuclear antigen (PCNA), vinculin and epidermal growth factor receptor (EGFR), as well as the intracellular calcium (Ca^2+^) signaling.

## Materials and methods

### Reagents

Ethanol (>98%), *n*-hexane (99%), dichloromethane (99.5%), ethyl acetate (99.5%), *n*-butanol (≥99.5%), methanol (≥99.8%), hydrochloric acid (37%), ammonium hydroxide (30%) and formic acid (98%) were purchased from Vetec Quimica Fina Ltda (Duque de Caxias, Rio de Janeiro, Brazil). Fluo-4/AM was purchased from Invitrogen (Eugene, USA). Enhanced chemiluminescence (ECL-plus Western Blotting Detection System) and peroxidase conjugated antibodies were purchased from Amersham Biosciences (Buckinghamshire, UK). All other reagents and standards were purchased from Sigma Aldrich Chemical Co. (St. Louis, MO, USA). Milli-Q Academic Water Purification System (Millipore Corp., Billerica, MA) was used to obatin the ionexchanged water. All reagents were of analytical grade.

### Plant material and alkaloid and acetogenin-rich fraction

*A. crassiflora* fruits were collected in the northern region of Minas Gerais State, Brazil, in March 2017. Voucher specimens (HUFU68467) were deposited in the herbarium of the Federal University of Uberlandia. The peels were quickly removed from the fresh fruits and crushed, and the obtained powder was stored at -20 °C until the moment of extraction. The dried and powdered peels (1.0 kg) were extracted for three days by maceration with 6 L of 98% ethanol at 25 °C. After filtration, ethanol was removed under reduced pressure using a rotary evaporator (Bunchi Rotavapor R-210, Switzerland) at 40 °C. This process was repeated until the last extract turned colorless (54.2 g, 5.42%). The alkaloid and acetogenin-rich fraction (AF-Ac) was obtained by a liquid-liquid extraction [10]. Briefly, the ethanol extract (10.0 g) was diluted in methanol:water (9:1, v/v, 200 mL), filtered and extracted using *n*-hexane (4 × 200 mL, 0.17 g), dichloromethane (4 × 200 mL, 0.31 g), ethyl acetate (4 × 200 mL, 2.71 g) and *n*-butanol (4 × 200 mL, 2.65 g). Additionally, an aqueous fraction (0.61 g) was obtained. All the phases were concentrated under reduced pressure at 40 °C, frozen and lyophilized (L101, Liobras, SP, Brazil). To confirm the presence of alkaloids, the fractions were analyzed by thin layer chromatography (TLC) (S1 Fig). The alkaloids and acetogenins were concentrated in the dichloromethane fraction. The resulting alkaloid and acetogenin-rich fraction was maintained at -20°C until use.

### Ultra-High-Performance Liquid Chromatography - Electrospray Ionization-tandem Mass Spectrometry (UHPLC-ESI/MS^n^)

The UHPLC-ESI/MS^n^ analysis of AF-Ac was done on an Agilent Q-TOF (model 6520) apparatus (Agilent, Santa Clara, CA, USA), operating in the positive mode. Methanol:water (4:1) was used as solvent system and the AF-Ac infused at the source at 200 µL/h. The parameters of chromatography were: Agilent Zorbax model 50 x 2.1 mm column, particles of 1.8 µm and pore diameter of 110 Å, mobile phase: water (0.1% formic acid, v/v) (A) and methanol (B). The gradient solvent system for B was: 2% (0 min); 98% (0-15 min); 100% (15-17 min); 2% (17-18 min); 2% (18-22 min), 0.35 mL/min and detection at 280 and 360 nm. The parameters of ionization were: 58 psi nebulizer pressure, 8 L/min N_2_ at 220 °C, and 4.5 kVa energy in the capillary. Sequential mass spectrometry (MS/MS) analyses were done with different collision energies (5-30 eV). The peaks and spectra were processed using the Agilent’s MassHunter Qualitative Analysis (B07.00) software and tentatively identified by comparing its retention time (Rt), error values (ppm) and mass spectrum with reported data [18].

### Cell culture

Human hepatocellular carcinoma cell line HepG2 was obtained from the American Type Culture Collection (ATCC HB-8065). HepG2 cells were cultured at 37°C in 5% CO_2_ in DMEM (GIBCOTM, Invitrogen Corp., Carlsbad, CA) supplemented with 10% fetal bovine serum (FBS), 4.5 g/L glucose, 1 mM sodium pyruvate, 50 units/mL penicillin, and 50 mg/mL streptomycin. Prior to addition of the treatments, cells were grown to 80-90% confluency and synchronized by incubating in serum-free medium (100% DMEM) for 24 h. The human peripheral blood mononuclear cells (PBMC) were purified using Histopaque-1077. All experimental procedures were carried out in accordance with the Code of Ethics of the World Medical Association (Declaration of Helsinki) and were approved by the Institutional Review Board of the Federal University of Uberlandia (no. 1.908.151) The informed consent was obtained from all subjects. Briefly, in conical tube 3 mL of EDTA-anticoagulated whole blood from three healthy volunteers was carefully layered onto 3 mL of Histopaque-1077 and then centrifuged at 400 *xg* for 30 min. PBMC were collected in plasma/Hitopaque-1077 interface and washed with 10 mL of Hank’s Balanced Salt Solution without calcium. Cells were suspended in RPMI-1640 supplemented with 10% of fetal bovine serum (Gibco), 2 mM L-glutamine, 100 U/mL penicillin and 100 µg/mL streptomycin. Semi-confluent (80% to 90%) cell cultures were used in all studies. The HepG2 cells were plated and then, 24 h later the AF-Ac treatment was done.The cells were then incubated with various concentrations (0-500 µg/mL) of AF-Ac for 24 and/or 48 h. Control group consisted of cells without addition of AF-Ac incubated only with vehicle (medium containing 0.05% DMSO). After 24 and/or 48 h, the cells and medium were collected. Protein contents in cells and medium were quantified by Bradford method [19].

### Cell viability

HepG2 and PBMC cells were seeded in 96-well microplate at 0.2 × 10^6^ cells/well and treated with AF-Ac (diluted in DMEM medium containing 0.05% DMSO for HepG2 cells or diluted in RPMI-1640 medium containing 0.05% DMSO for PBMC cells) or vehicle (control, DMEM medium containing 0.05% DMSO for HepG2 cells; RPMI-1640 medium containing 0.05% DMSO for PBMC cells) for 24 h. Then, 100 µL of 5 mg/mL (3-(4,5-dimethylthiazolyl-2)−2,5-diphenyltetrazolium bromide) solution was incubated with the supernatant at 37 °C for 2 h in 5% CO_2_. Next, dimethyl sulfoxide (DMSO) was added and the cell viability was analyzed by absorbance of the purple formazan from viable cells at 570 nm (Molecular Devices, Menlo Park, CA, USA).

### Cellular proliferation assay

HepG2 cells were grown in 24 well-plates. FBS was removed for overnight and then the cells were treated with 50 µg/mL AF-Ac or vehicle (control, DMEM medium containing 0.05% DMSO). *In vitro* cell proliferation assay was assessed by manual counting in Neubauer chamber using optic microscopy at 6, 12, 24 and 48 h, as previously described [20].

### Migration assay

HepG2 cells were grown in 12-well plates and treated with 50 µg/mL AF-Ac or vehicle (control, DMEM medium containing 0.05% DMSO) for 48 h. Migration assay was performed as previously described [21]. The wound was achieved by scratching a pipette tip across the cell monolayer (approximately 1.3 mm in width); 1 µM hydroxyurea was added to prevent the proliferation [22]. The wound area was measured using the Northern Eclipse (Empix, Mississauga, Canada) software, and the percentage of wound closure at each time point was derived by the formula: (1 –[current wound size/initial wound size]) × 100.

### Western blot analyses

HepG2 cell lysates in SDS-sample buffer containing an additional 100 mM Tris-HCl pH 8.0 and 25% glycerol were boiled for 5 min and equal amounts of total protein (25 µg/mL) were separated by 12% SDS-PAGE gel. To better take advantage of the western blot, in which triplicates of each sample are present, the whole membranes were cut into strips for the different antibodies tested. The blots were cut prior to hybridization with antibodies. Images of all blots as they are, and all replicates performed are shown in S16 Fig. For protein detection, specific primary antibodies against proliferating cell nuclear antigen PCNA (mouse, 1:1,000;), vinculin (mouse, 1:1,000, Cell Signaling Technology), epidermal growth factor receptor EGFR (mouse, 1:1,000 Santa Cruz Biotechnology, Dallas, TX) and β-actin (mouse, 1:1,000; Santa Cruz Biotechnology, Dallas, TX) were used. The primary antibody incubation proceeded for 2 h at room temperature. After being washed, blots were incubated with horseradish peroxidase-conjugated specific secondary antibody (anti-mouse or anti-rabbit, 1:5,000; Sigma-Aldrich) at room temperature for 1 h. Immune detection was carried out using enhanced chemiluminescence (ECL plus; Amersham Biosciences) [23]. Western blot digital images (8-bit) were used for densitometric analysis using ImageJ (National Institutes of Health, Bethesda, MD).

### Immunofluorescence

Confocal microscopy examination of immunofluorescence in HepG2 cells was performed as described [24]. Cells were seeded onto 6-well culture dishes and incubated with 50 µg/mL AF-Ac or vehicle (control, DMEM medium containing 0.05% DMSO) for 24 h. Then, cells were fixed with 4% paraformaldehyde, permeabilized with PBS 1X/Triton 0.5% and blocked with PBS (10% BSA, 0.5% Triton 0.5% and 5% goat serum) for 1 h. Cells were incubated with anti-EGFR antibody (anti-mouse, 1:100; Abcam, MA, USA) for 2 h at room temperature, followed by incubation with anti-mouse secondary antibody conjugated with Alexa 488 (1:500; Life Technologies) for 1 h. Isotype control was used to assess non-specific binding under the same experimental conditions. Images were obtained using a Zeiss LSM 510 confocal microscope (Thornwood, NY, USA) equipped with a 63×/1.4 NA objective with excitation laser at 488 nm and emission bandpass filter at 505-550 nm.

### Detection of Ca^2+^ signals

Intracellular Ca^2+^ was monitored in individual cells by time lapse confocal microscopy, as described previously [25]. Briefly, HepG2 cells were incubated with Fluo-4/AM (6 μM) for 30 min at 37 °C in 5% CO_2_ in HEPES buffer with or without 10 mM EGTA. Then, coverslips containing cells were transferred to a perfusion chamber on the stage of the Zeiss LSM510 confocal imaging system equipped with a Kr-Ar laser. Nuclear and cytosolic Ca^2+^ signals were monitored in individual cells during stimulation with 50 µg/mL AF-Ac using a ×63, 1.4 NA objective lens. Fluo-4/AM was excited at 488 nm and observed at 505-550 nm. Changes in fluorescence were normalized by the initial fluorescence (F0) and were expressed as (F/F0) × 100. During the 600 s for the calcium signalling experiments, the cells were perfused with HEPES solution without fetal bovine serum, grown factor and molecules that can themselves alter the calcium signalling.

### Statistical analysis

The graphics and statistical analyzes were done using SigmaPlot (Systat Software, Point Richmond, USA) and Prism (GraphPad Software, San Diego, USA). The data were expressed as mean ± SD and the statistical significance was tested using Student’s t test, one-way or two-way ANOVA followed by Dunnett or Bonferroni test. *p* value < 0.05 was taken to indicate statistical significance.

## Results

### Identification of alkaloids and acetogenins by Ultra-High-Performance Liquid Chromatography-Electrospray Ionization-tandem Mass Spectrometry (UHPLC-ESI/MS^n^)

The alkaloid and acetogenin profile of AF-Ac was performed by UHPLC-ESI–MS^n^. The presence of ions *m/z* attributed to alkaloids and acetogenins was confirmed in the positive mode by high resolution “zoom scan” analysis. Isopiline, isoboldine, isocorydine, anonaine, nuciferine, xylopine, stephalagine, liriodenine and atherospermidine were the alkaloids found in AF-Ac [11, 26–31], whereas bullatanocin, bullatacin/squamocin, annomontacin and desacetyluvaricin/ isodesacetyluvaricin were the acetogenins found in AF-Ac [32] (Fig 1 and Table 1). The chemical structures of the alkaloids and acetogenins identified in the AF-Ac fraction are shown in Figs 2 and 3, respectively. The sequential mass spectra can be found as supplementary material online (S2-S14 Fig).

**Fig 1.**
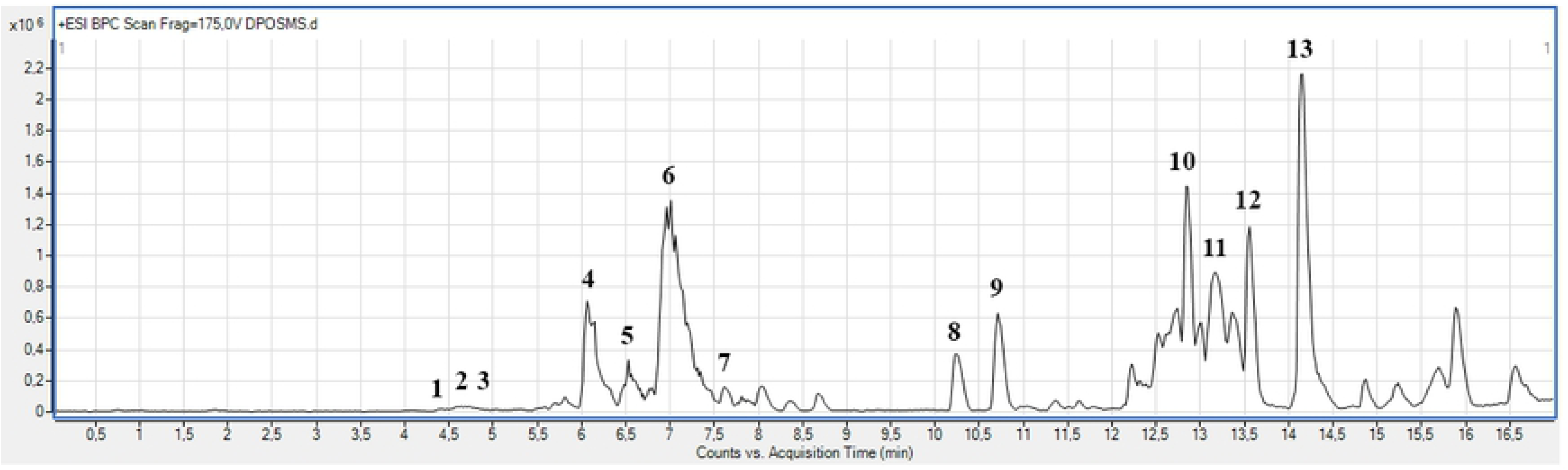
**Chromatogram of the alkaloid and acetogenin-rich fraction from *Annona crassiflora* fruit peel (AF-Ac) by HPLC-ESI-MS/MS (positive mode).**

**Fig 2.**
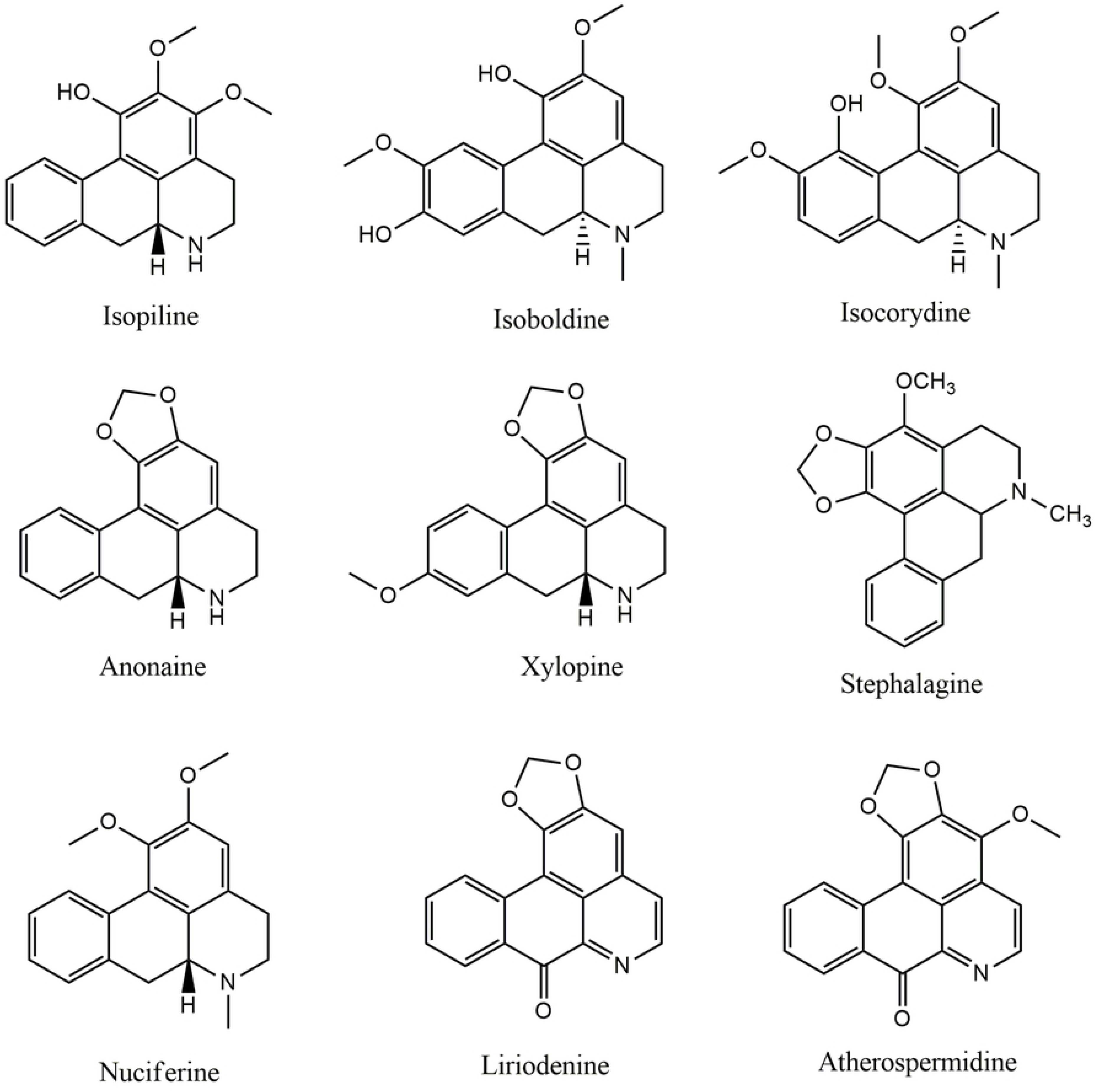
**Chemical structures of the aporphine alkaloids identified in the AF-Ac fraction from *A. crassiflora* fruit peel.**

**Fig 3.**
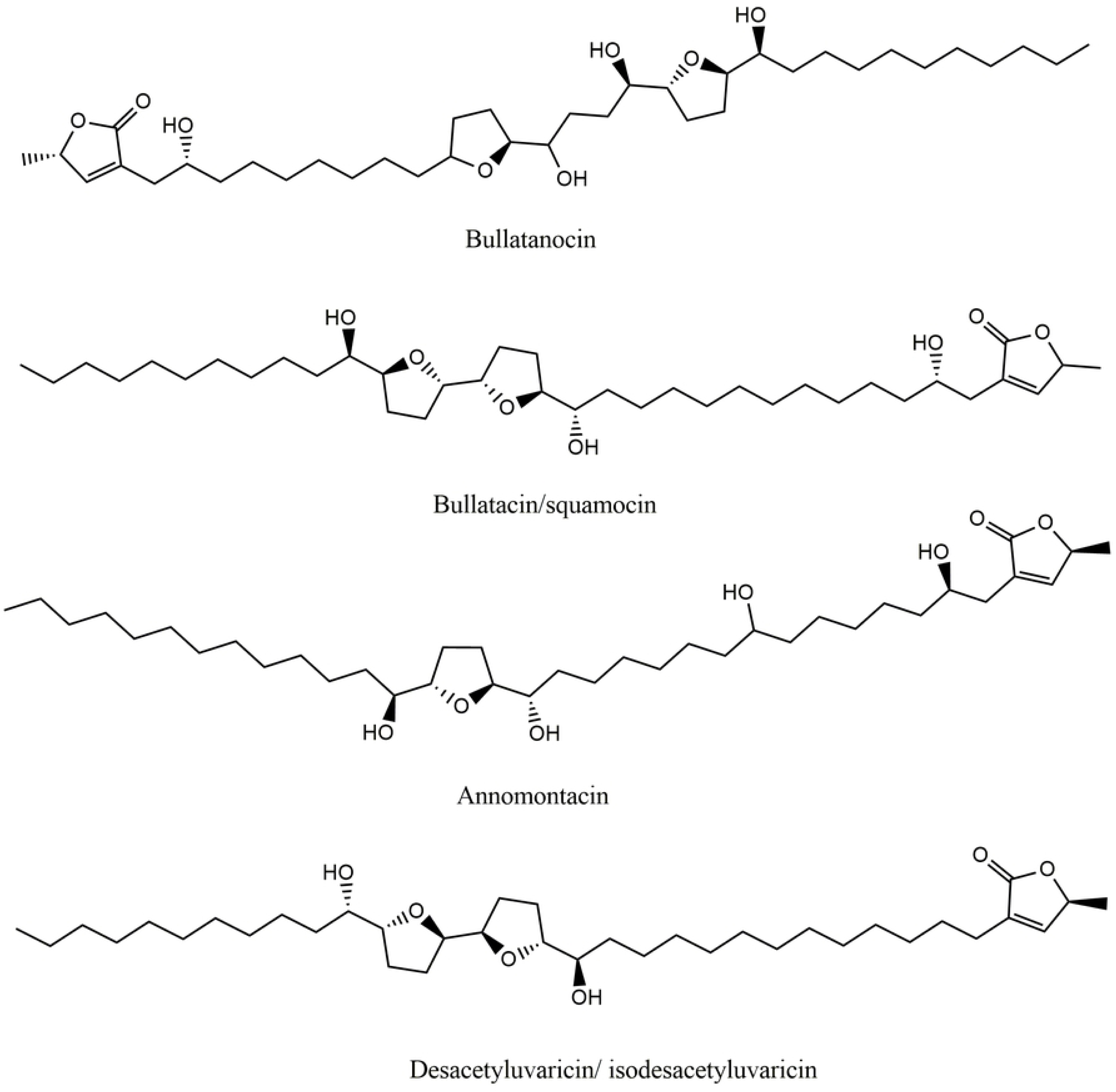
**Chemical structures of the acetogenins identified in the AF-Ac fraction from *A. crassiflora* fruit peel.**

**Table 1.**
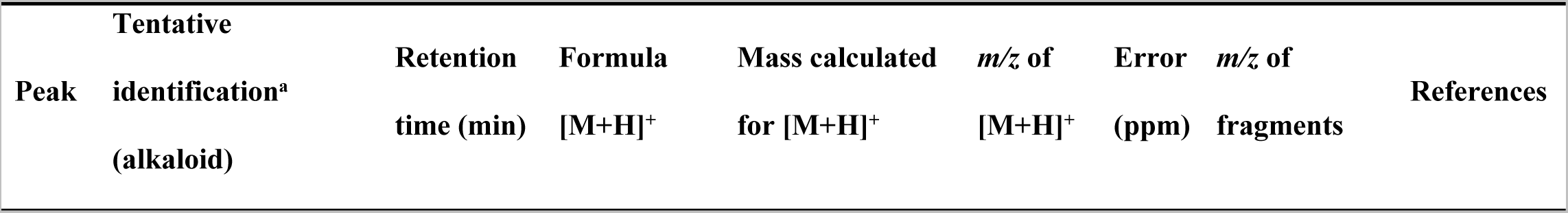
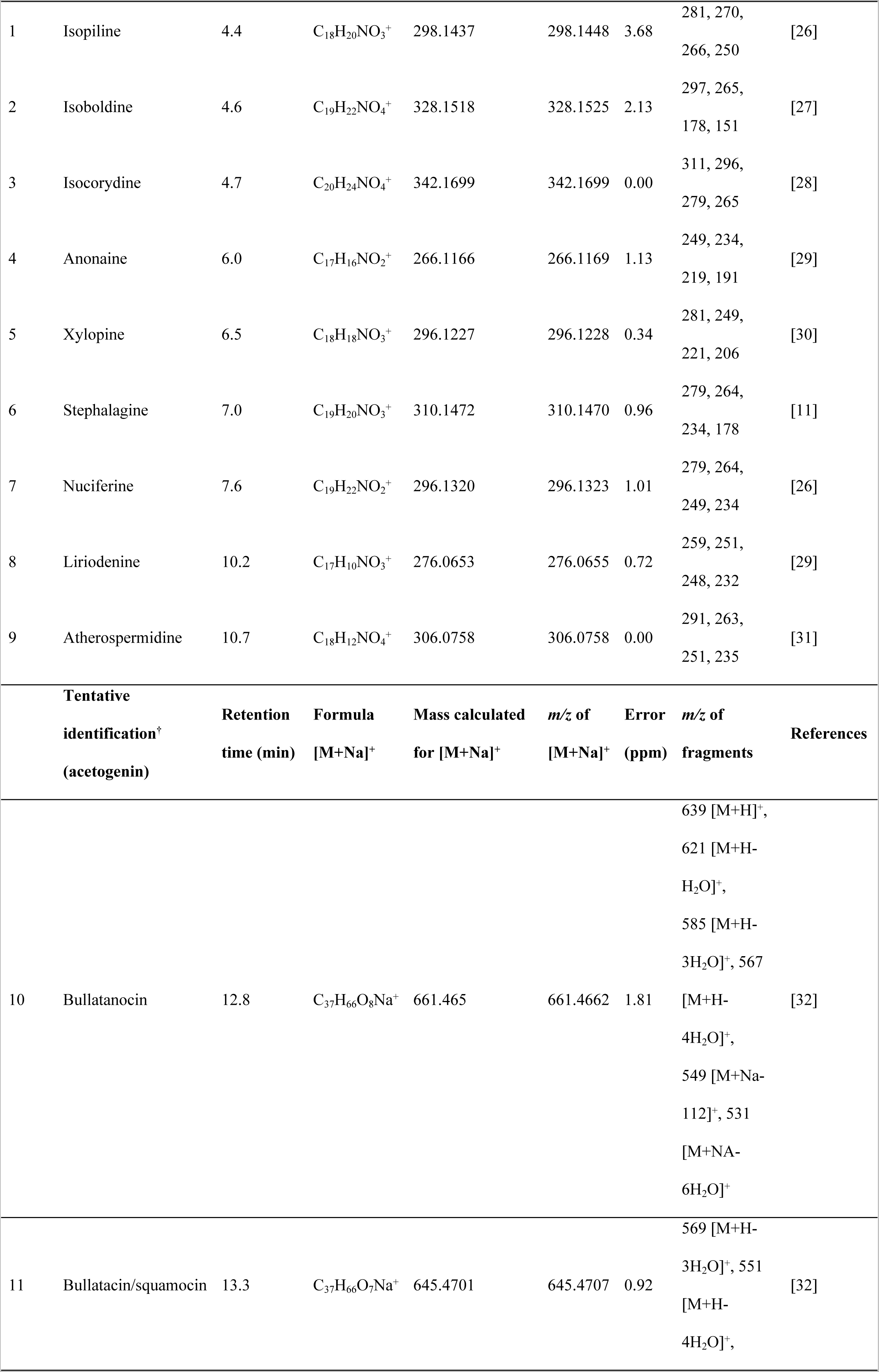
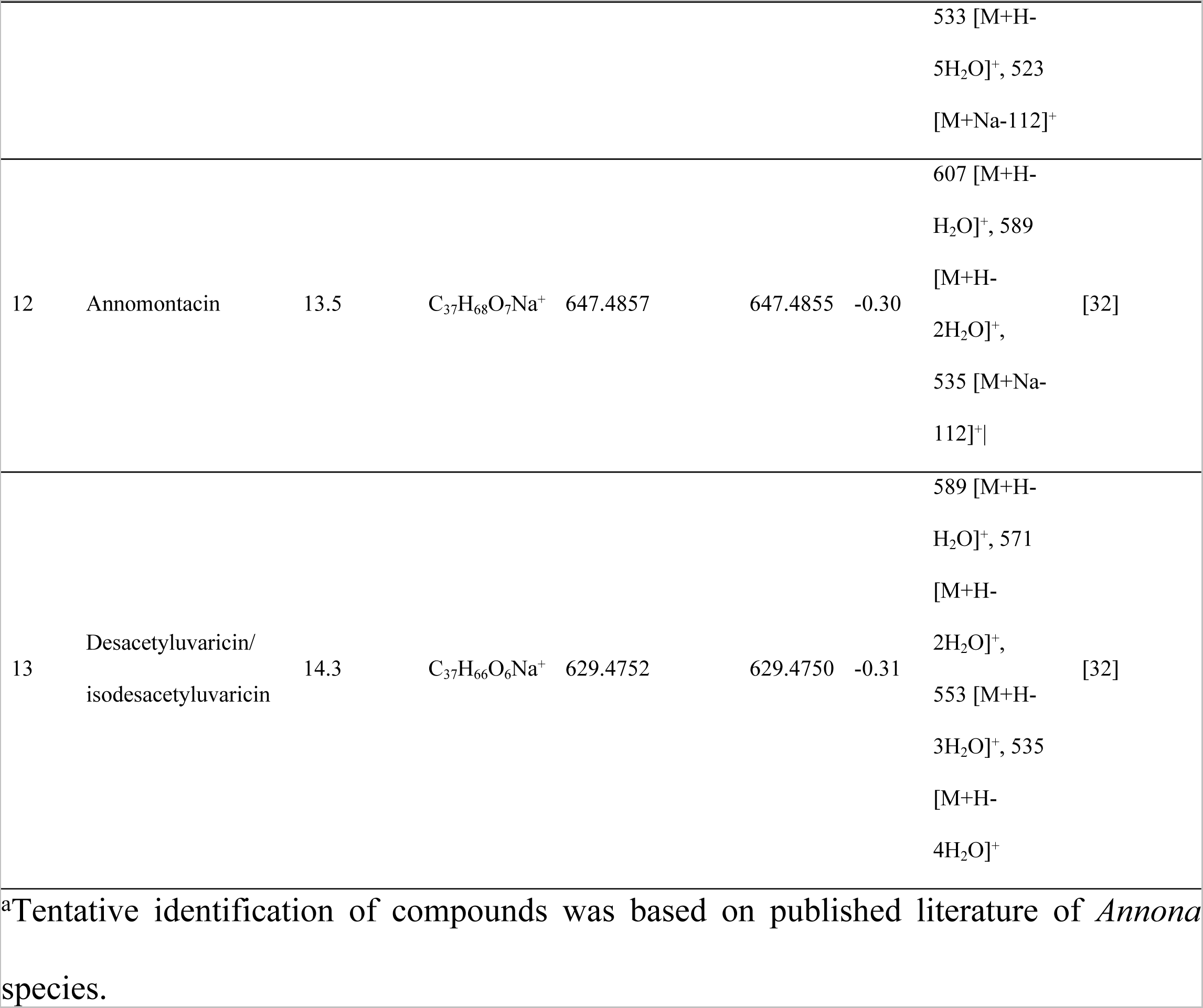
Alkaloids and acetogenins identified in the alkaloid and acetogenin-rich fraction from *Annona crassiflora* fruit peel (AF-Ac) by UHPLC-ESI/MS^n^ (positive mode).

### AF-Ac reduces HepG2 cell viability

Fig 4 shows cell viability of HepG2 and PBMC cells treated with different concentrations of AF-Ac for 24 h. AF-Ac was able to reduce HepG2 cell viability at 50, 250 and 500 µg/mL, compared to untreated control cells (cells treated only with vehicle) (25.7 ± 2.8, 77.0 ± 1.8 and 83.3 ± 1.3% of reductions, respectively, *p* < 0.001) (Fig 4A). However, AF-Ac was cytotoxicity for PBMC cells only at 500 µg/mL (Fig 4B). As was observed with PBMC cells, the AF-Ac fraction at a concentration of 50 µg/mL did not affect the cell viability of fibroblasts treated for 24 h (S15 Fig).

**Fig 4.**
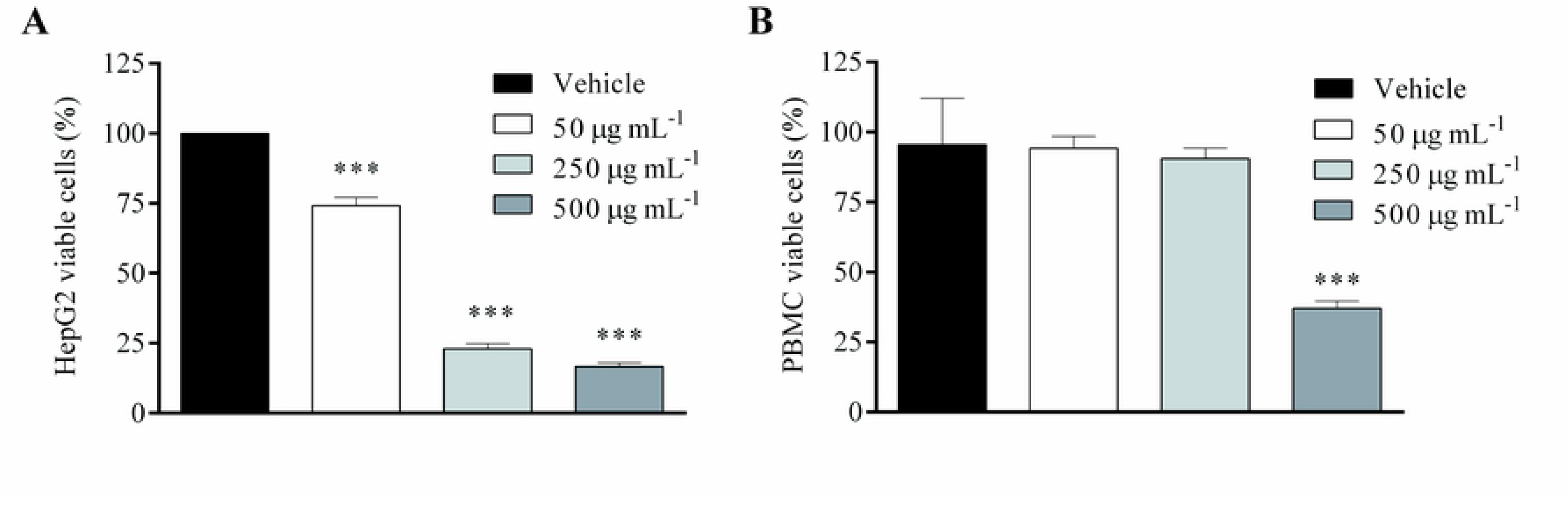
**Cell viability of HepG2 (A) and PBMC (B) cells treated with the alkaloid and acetogenin-rich fraction of *Annona crassiflora* fruit peel (AF-Ac) or vehicle (control, cells treated with DMEM medium containing 0.05% DMSO for HepG2 cells or RPMI-1640 containing 0.05% DMSO for PBMC cells). Results (mean ± SD, n = 3) expressed as the percentage of viable cells compared to the vehicle group. Significance levels are indicated by \*\*\**p* < 0.001 when compared to control (one-way ANOVA and Dunnett as posttest).**

### AF-Ac reduces HepG2 cell proliferation

We investigated whether AF-Ac presents antiproliferative effect in HepG2 cells since it reduced its viability. Incubation of HepG2 cells with 50 µg/mL AF-Ac for 48 h led to a reduction in cell proliferation (75.2 ± 10.5%, *p* < 0.001) (Fig 5A). This result was similar to cells in medium with 0% fetal bovine serum. After 24 h incubation with AF-Ac, the expression of PCNA, a marker of cell proliferation, in HepG2 cells was analyzed by Western blotting. In accordance with the cell proliferation assay, AF-Ac at the dose of 50 µg/mL decreased PCNA expression (*p* < 0.05) (Figs 5B and 5C).

**Fig 5.**
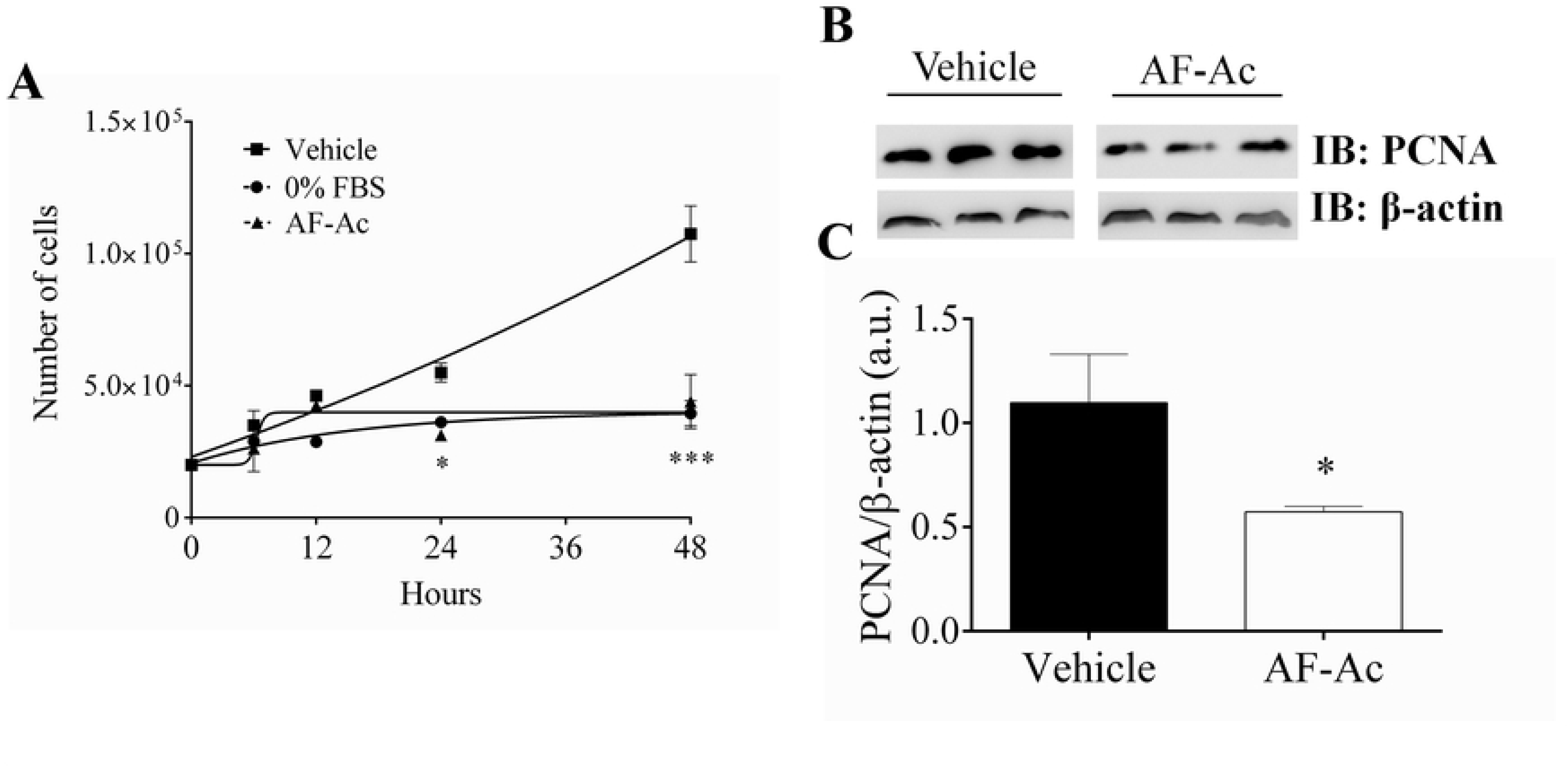
**Cell growth assay of HepG2 cells at 6, 12, 24 and 48 h after stimulation with 50 µg/mL AF-Ac or vehicle (control, cells treated with DMEM medium containing 0.05% DMSO), triplicate in 3 individual experiments (A). Representative immunoblotting of total HepG2 cell lysates probed with anti-PCNA and anti-β-actin, used as protein loading control (B). Immunoblotting densitometry analysis. Results show β-actin normalized proteins expression (n = 3 individual experiments/group) (C). Values are expressed as mean ± SD. Significance levels are indicated by \**p* < 0.05 (unpaired t-test) and \*\*\**p* < 0.001 (two-way ANOVA followed by Bonferroni’s post hoc test) when compared to the vehicle group. Each protein was analyzed in cropped membranes of different Western blots along with other proteins. Full-length blots are presented in S16 Fig.**

### AF-Ac reduces HepG2 cell migration

Following these observations, we investigated the influence of AF-Ac on the migration of HepG2 cells. Thus, a scratch assay was made in the presence or absence of AF-Ac (Fig 6A). After 48 h, AF-Ac decreased the healing process (47.1 ± 1.5% of healing) when compared with untreated cells (74.2 ± 3.5% of healing) (*p* < 0.001) (Fig 6B). In order to check if the reduced healing observed during stimulation with AF-Ac is due to alterations on the focal adhesion points, we performed immunoblotting for vinculin. However, vinculin levels were not affected in HepG2 cells treated with 50 µg/mL AF-Ac (Figs 6C and 6D).

**Fig 6.**
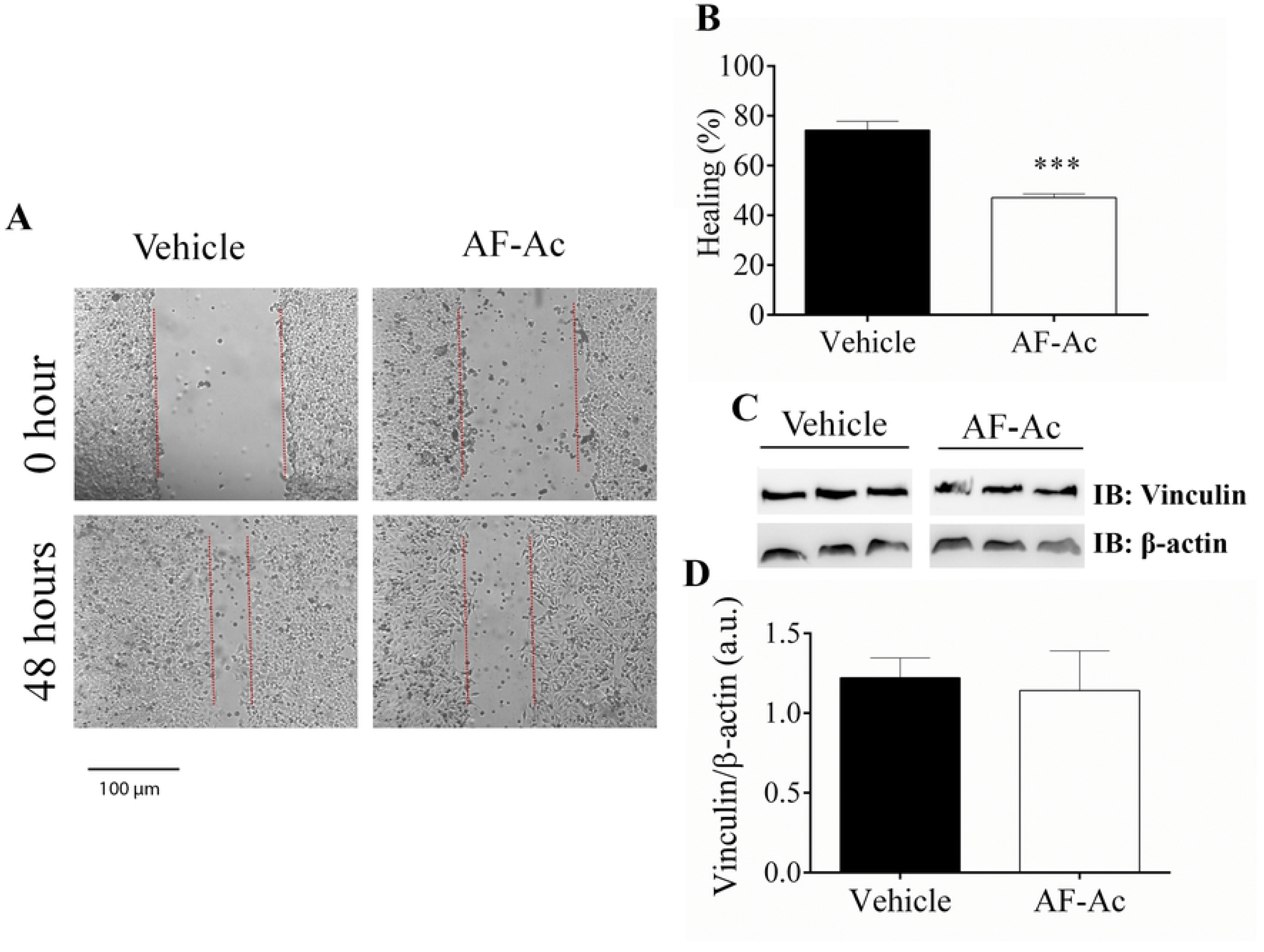
**Representative image of *in vitro* wound healing assay performed with HepG2 cells. Images were selected from a representative well 48 h after stimulation with 50 µg/mL AF-Ac or vehicle (control, cells treated with DMEM medium containing 0.05% DMSO). Scale bar = 100 μm (A). Average of wound healing closure 48 h after stimulation with AF-Ac (n = 5 wells/group, for each time point). Results represent % of initial wound area (0 h) (B). Representative cropped immunoblotting of total HepG2 cell lysates probed with anti-vinculin and anti-β-actin, used as protein loading control (C). Immunoblotting densitometry analysis. Results show β-actin normalized proteins expression (n = 3 individual experiments/group) (D). Values are expressed as mean ± SD. Significance levels are indicated by \*\*\**p* < 0.001 (unpaired t-test) when compared to the vehicle group. Each protein was analyzed in cropped membranes of different Western blots along with other proteins. Full-length blots are presented in S16 Fig.**

### Ac reduces EGFR in HepG2 cells

EGFR is known to play an important role in the regulation of cell proliferation in HepG2 cells. As revealed by immunofluorescence assay (Figs 7A and 7B), EFGR levels were reduced in HepG2 cells treated with 50 µg/mL AF-Ac (*p* < 0.05). This result is in accordance with data obtained by immunoblotting assay, with HepG2 cells treated with 50 µg/mL AF-Ac presenting decreased expression of EGFR (*p* < 0.01) (Figs 7C and 7D).

**Fig 7.**
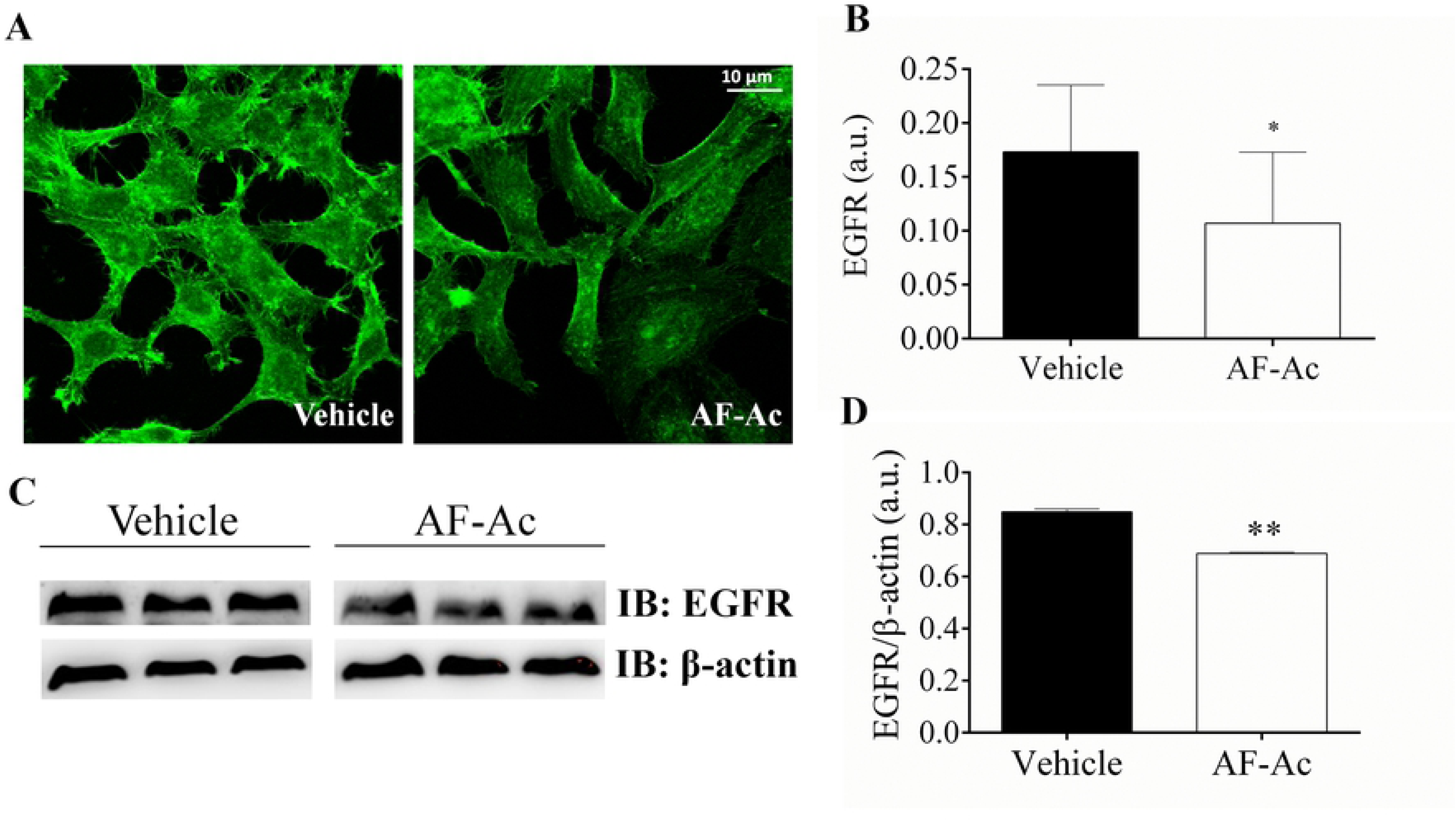
**Representative immunofluorescence images of HepG2 cells treated with 50 µg/mL AF-Ac or vehicle (control, cells treated with DMEM medium containing 0.05% DMSO) labeled with specific anti-EGFR (green) antibody. Scale bar = 10 μm (A). Average number of EGFR–positive regions for the cell types analyzed are shown as EGFR/1000 μm^2^ on each respective graph (n = 20 cells/group) (B). Representative cropped immunoblotting of total HepG2 cell lysates probed with anti-EGFR and anti-β-actin, used as protein loading control (C). Immunoblotting densitometry analysis. Results show β-actin normalized proteins expression (n = 3 individual experiments/group) (D). Values are expressed as mean ± SD. Significance levels are indicated by \*\**p* < 0.01 (unpaired t-test). Each protein was analyzed in cropped membranes of different Western blots along with other proteins. Full-length blots are presented in S16 Fig.**

### AF-Ac increases intracellular Ca^2+^ in HepG2 cells

HepG2 cells were loaded with fluo4/AM and assayed for Ca^2+^ signals during AF-Ac stimulation. Fig 8A shows that there was an increase in intracellular-free Ca^2+^ when HepG2 cells were exposed to 50 µg/mL AF-Ac. After 60 s intracellular Ca^2+^ rose to peak levels, as shown by the increase in fluorescence (Fig 8B). We also performed experiments in the absence of extracellular Ca^2+^ to explore the relative contribution of intracellular Ca^2+^ pools to the overall response induced by AF-Ac. Thus, 10 mM EGTA was added to the Ca^2+^-free HEPES buffer, chelating extracellular-free Ca^2+^ levels. AF-Ac triggered Ca^2+^ wave in HepG2 cells with a peak at 480 s after AF-Ac exposure (Fig 8C). No difference was observed between Ca^2+^ levels from nucleus and cytosol (Fig 8D).

**Fig 8.**
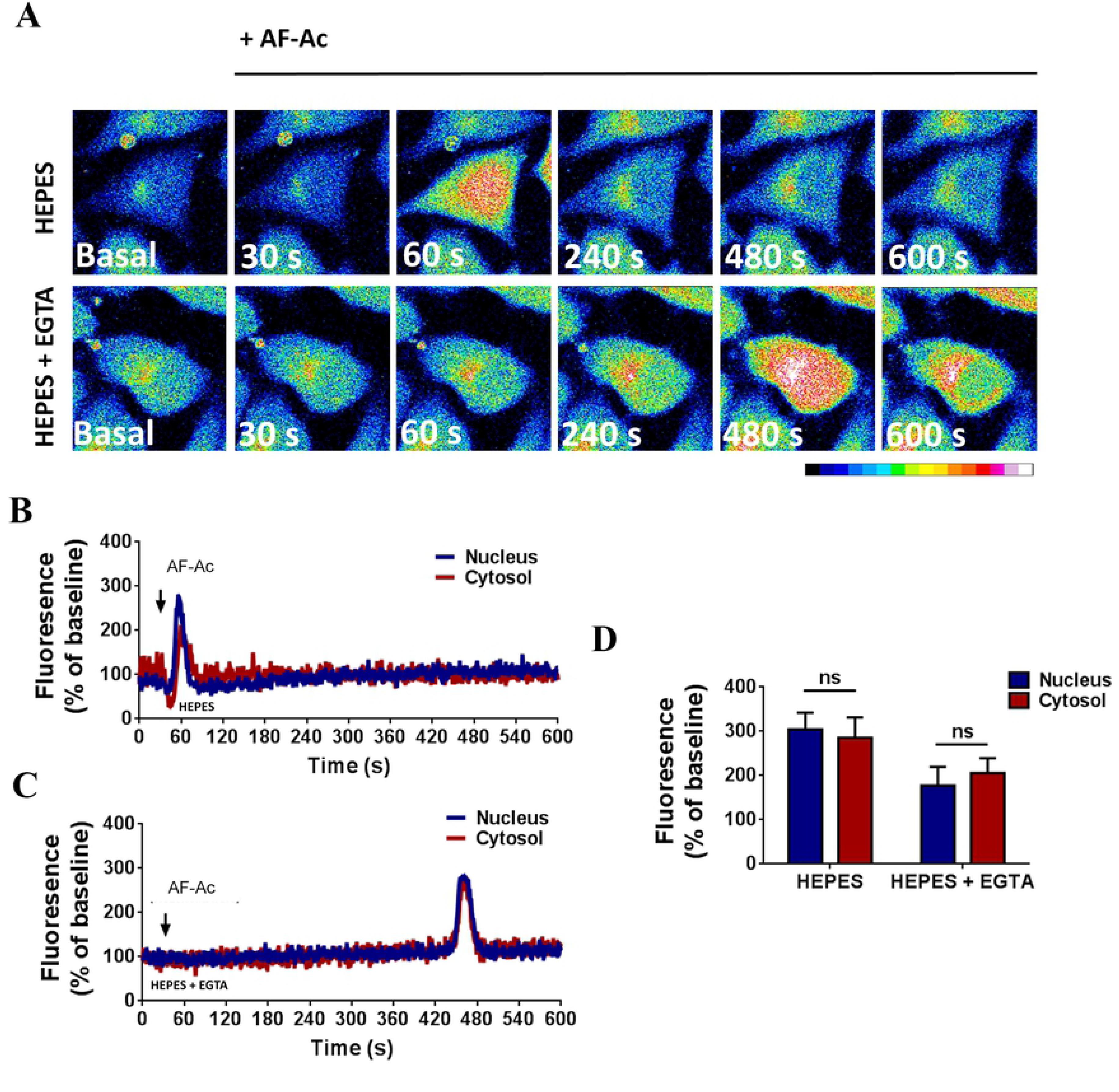
**Confocal serial images of HepG2 cells, loaded with fluo-4/AM and stimulated with 50 µg/mL AF-Ac for 30, 60, 240, 360, 480 and 600 s. Dashed yellow regions represent the nuclear region. Images were pseudocolored according to the scale shown at the bottom. Scale bar = 10 μM (A). Representative time course of nuclear and cytosol fluorescence levels of HepG2 cells, in the presence (B) or absent (C) of EGTA, stimulated with AF-Ac. Black arrow indicates initial AF-Ac stimulation and fluorescence level is expressed as % of basal fluorescence. Average nuclear and cytosol fluorescence peaks (n = 20 cells) of each cell group and condition throughout the time-course; fluorescence level is expressed as % of basal fluorescence (D). Values are expressed as mean ± SD. ns = *p* > 0.05 (unpaired t-test).**

## Discussion

The search for natural agents capable of controlling tumor growth and presenting low toxicity on normal healthy cells has gained prominence in the treatment of cancer [3, 4, 33, 34]. Whole plants or herbal extracts/fractions have been used rather than isolated molecules due to their more affordable access and synergistic interaction between the compounds that may increase the biological effects [35]. Numerous alkaloids from medicinal plants and herbs have showed antiproliferative and anticancer effects on a wide category of cancers both *in vitro* and *in vivo* [4]. Another example is the annonaceous acetogenins, which have been identified as cancer growth inhibitors and/or apoptotic agents [9]. In the present study, we showed the anticancer potential of an alkaloid and acetogenin-rich fraction from *A. crassiflora* fruit peel, named here as AF-Ac, by evaluating its antiproliferative properties in human liver carcinoma cells (HepG2) *in vitro*.

First, we performed an ethanolic extraction of the fruit peel of *A. crassiflora* and followed this with a liquid-liquid fractionation of the crude extract to obtain an alkaloid and acetogenin-rich fraction. Our findings indicated that the dichloromethane fraction had alkaloids and acetogenins, which was confirmed by UHPLC-ESI/MS^n^ and TLC analyses (supplementary material). Interestingly, all the alkaloids found in AF-Ac are aporphine alkaloids, such as annonaine, isopoline, isoboldine, isocorydine, liriodenine, stephalagine, nuciferine, atherospermidine and xylopine. Until now, stephalagine was the only alkaloid isolated and characterized in *A. crassiflora* fruit peel [11]. The retention time, exact mass and MS/MS spectra of stephalagine showed in the present study corroborates the data reported by Justino, Barbosa (10). In addition, acetogenins such as bullatanocin, bullatacin/squamocin, annomontacin and desacetyluvaricin were identified for the first time in the fruit peel of *A. crassiflora*.

Annonaceous acetogenins, more specifically bis-tetrahydrofuranic (THF) acetogenins like that found in AF-Ac, have shown cytotoxicity in HepG2 cells through the induction of cell-cycle arrest and induction of the apoptotic mitochondrial pathway involving complexation with Ca^2+^ [36–38]. Previous studies have also reported that aporphine alkaloids, such as liriodenine, have prominent cytotoxic effects in several cancer cell lines such as inducing G1 cell cycle arrest and repressing DNA synthesis in HepG2 cells, and reducing cell growth and inducing apoptosis in human breast cancer MCF-7 cells through inhibition of Bcl02, cyclin D1 and vascular endothelial growth factor [6, 39]. Anonaine also showed cytotoxic effects in HepG2 cells and caused DNA damage associated with increased intracellular nitric oxide and ROS, glutathione depletion, disruptive mitochondrial transmembrane potential and activation of caspases 3, 7, 8 and 9 [40]. Moreover, anonaine also up-regulated the p53 and Bax expression [40]. Isocorydine decreased the viability of hepatocellular carcinoma (HCC) and HepG2 cells [41, 42]. Nuciferine is considered as an anti-tumor agent against human neuroblastoma and mouse colorectal cancer *in vitro* and *in vivo*, through inhibiting the PI3K-AKT signaling pathways [43]. Furthermore, a study done by Kang, Lee (44) showed that nuciferine inhibited the growth of breast cancer cells. Xylopine and isoboldine, two other aporphine alkaloids found in AF-Ac, were cytotoxic to HepG2 cells and were able to arrest G2/M cycle [45, 46].

AF-Ac at the dose of 50 µg/mL effectively decreased HepG2 cell viability and reduced cell proliferation. It is worth mentioning that the AF-Ac at 50 µg/mL was not cytotoxic for PBMC and fibroblast cells. The objective of using PBMC and fibroblast cells as controls is to demonstrate that the AF-Ac fraction is not cytotoxic to healthy cells, since these human non-cancer cells are potentially useful models for cell viability testing using plant extracts [47–50]. Also, PBMC cells represent the whole metabolic status and an excellent model for assessing the differences or changes associated with pathophysiological conditions [47, 51]. Additionally, a study conducted by our research group also showed no cytotoxicity of the AF-Ac fraction in Vero cells [11].

Consistent with the results of MTT and proliferation assays, AF-Ac decreased the expression of PCNA in the HepG2 cells. PCNA is a cell nuclear protein whose expression is correlated with DNA replication, regulating the transition from G1 phase to S phase, and is connected with the proliferation of tumor cells [52]. The antiproliferative potential of some alkaloids and acetogenins has been associated with their capacity to reduce PCNA expression. A study done by Long and Li (53) showed the reduction of PCNA expression by an alkaloidal fraction from aerial parts of *Oxytropis ochrocephala* in mice hepatocellular carcinoma. In addition, the antitumor potential of berberine and matrine was demonstrated in the inhibition of the expression of PCNA in ovarian cancer and lung adenocarcinoma cells, respectively [54, 55]. Annomuricin E, an acetogenin isolated from *A. muricata* leaf, was also able to down-regulate the PCNA expression in HT-29 colon cancer cells [56].

As well as the PCNA, EGFR also plays an important role in the regulation of cell proliferation [57]. EGFR overexpression might contribute to deregulated cellular processes, such as uncontrolled proliferation, invasion, DNA synthesis, angiogenesis, cell motility and inhibition of apoptosis, which makes it a molecular target for tumor therapy [57]. In the present study, AF-Ac reduced the expression of EGFR in the HepG2 cells, as showed by Western blot and immunofluorescence assays. Isocorydine, an aporphine alkaloid found in AF-Ac, has previously shown cytotoxic effects in HepG2 cells and, by a docking analysis, and has inhibitory activity against EGFR [42]. Dicentrine, another aporphine alkaloid, has been shown to exert cytotoxic activity towards cancer cells by binding to EGFR [7, 33]. In addition, acetogenins influences EGFR signaling to induce cell cycle arrest and inhibit cytotoxic cell survival [58]. These findings indicate that EGFR and PCNA signaling pathways might play a role in mediating the antiproliferative activity of AF-Ac on HepG2 cells.

Studies have demonstrated that alkaloids and acetogenins may inhibit cell migration and metastasis of cancer cells [59–61]. Here, we showed the capacity of AF-Ac to decrease cell migration of HepG2 cells without vinculin overexpression. The capacity of tumor cells to migrate is essential for many physiological processes including tumor invasion, angiogenesis and metastasis [62]. The filamentous (F)-actin-binding protein vinculin is required for cell polarization and migration, having a key role on the formation of focal adhesion points [63]. Thus, cells with reduced expression of vinculin become less adherent and more motile [64]. Thus, AF-Ac might be acting on other targets involved with cell migration than vinculin expression.

Finally we investigated whether AF-Ac alters intracellular Ca^2+^ in the HepG2 cells, since nuclear Ca^2+^ was previously found to negatively regulate cell motility, invasion and proliferation [22, 65]. Of note, confocal analysis showed that AF-Ac increased intracellular Ca^2+^ through a process that involved Ca^2+^ influx which requires external calcium, when Ca^2+^ was present in the external media. In the absence of external Ca^2+^, exposure of HepG2 cells to AF-Ac also mobilized intracellular Ca^2+^. This suggests that the alkaloid and acetogenin-rich fraction induces a mobilization of intracellular Ca^2+^ stores. Ca^2+^ is a ubiquitous second messenger that regulates a wide range of activities in cells, such as secretion, contraction, metabolism, gene transcription, apoptosis and proliferation [66]. Studies have showed that nuclear Ca^2+^ buffering reduced cell proliferation in hepatocellular carcinoma cells by stopping cell cycle progression, modulating the promoter region activity of genes involved in cell proliferation and/or preventing the upregulation of the tyrosine kinase receptor [20, 24, 67]. In addition, nuclear Ca^2+^ buffering may turn cells more rigid and less motile due to the reduction of membrane fluctuations [22].

Although studies have showed that the mentioned aporphine alkaloids reduce Ca^2+^ influx [10, 68, 69], AF-Ac induced increases in cytosolic and nuclear Ca^2+^ in the HepG2 cells. It is well known that acetogenins induce an increase of cytosolic and mitochondrial Ca^2+^ in several cancer cells [38], which might explain the intracellular Ca^2+^ increase observed in the HepG2 treated with AF-Ac. The mechanism underlying the cytotoxicity and antiproliferative effect of acetogenins is modulated by the chelation of THF moieties with Ca^2+^ to form hydrophobic complexes, which may induce sustained increases in intracellular and mitochondrial Ca^2+^ concentrations, resulting in decrease of mitochondrial membrane potential that leads to the release of apoptotic initiators [38, 70]. Thus, the chelating ability of acetogenins with Ca^2+^ might contribute, at least in part, to the anti-migration and anti-proliferative capacities of AF-Ac in the HepG2 cells.

## Conclusions

In summary, our results have established the antiproliferative properties of AF-Ac on HepG2 cells and suggest that this effect is mediated, at least in part, by reducing PCNA and EGFR expression with a mobilization of intracellular Ca^2^. Although the biochemical mechanisms involved in the antiproliferative effect of the alkaloids and acetogenins from *A. crassiflora o*n HepG2 cells were not fully explored, this study is the first to identify the alkaloids and acetogenins present in the fruit peel of *A. crassiflora* and to demonstrate its antitumoral potential. Furthermore, the biological activities exercised by the AF-Ac fraction were observed in concentrations below the cytotoxic level. Thus, the use of this alkaloid and acetogenin-rich fraction in further *in vivo* assays is justified.

## Acknowledgments

The authors gratefully acknowledge the Institute of Biotechnology of the Federal University of Uberlândia and the Department of Physiology and Biophysics and the Liver Center of the Federal University of Minas Gerais for infrastructural support, Mário M. Martins and Paula Souza Santos for technical assistance in LC-MS analysis, and Luiz Ricardo Goulart for supervision.

## Supporting information

**S1 Fig. Analytical TLC (SiO2) of the alkaloid and acetogenin-rich fraction from *Annona crassiflora* fruit peel (AF-Ac) developed with CH_2_Cl_2_-MeOH-NH_4_OH (9:1:0.25) and reveled with IClPt reagent.**

**S2 Fig. HPLC-ESI-MS/MS of isopiline from the alkaloid and acetogenin-rich fraction from *Annona crassiflora* fruit peel (AF-Ac) (*m*/*z* 298 [M+H]^+^).**

**S3 Fig. HPLC-ESI-MS/MS of isoboldine from the alkaloid and acetogenin-rich fraction from *Annona crassiflora* fruit peel (AF-Ac) (*m*/*z* 328 [M+H]^+^).**

**S4 Fig. HPLC-ESI-MS/MS of isocorydine from the alkaloid and acetogenin-rich fraction from *Annona crassiflora* fruit peel (AF-Ac) (*m*/*z* 342 [M+H]^+^).**

**S5 Fig. HPLC-ESI-MS/MS of anonaine from the alkaloid and acetogenin-rich fraction from *Annona crassiflora* fruit peel (AF-Ac) (*m*/*z* 266 [M+H]^+^).**

**S6 Fig. HPLC-ESI-MS/MS of xylopine from the alkaloid and acetogenin-rich fraction from *Annona crassiflora* fruit peel (AF-Ac) (*m*/*z* 296 [M+H]^+^).**

**S7 Fig. HPLC-ESI-MS/MS of stephalagine from the alkaloid and acetogenin-rich fraction from *Annona crassiflora* fruit peel (AF-Ac) (*m*/*z* 310 [M+H]^+^).**

**S8 Fig. HPLC-ESI-MS/MS of nuciferine from the alkaloid and acetogenin-rich fraction from *Annona crassiflora* fruit peel (AF-Ac) (*m*/*z* 296 [M+H]^+^).**

**S9 Fig. HPLC-ESI-MS/MS of liriodenine from the alkaloid and acetogenin-rich fraction from *Annona crassiflora* fruit peel (AF-Ac) (*m*/*z* 276 [M+H]^+^).**

**S10 Fig. HPLC-ESI-MS/MS of atherospermidine from the alkaloid and acetogenin-rich fractionfrom *Annona crassiflora* fruit peel (AF-Ac) (*m*/*z* 306 [M+H]^+^).**

**S11 Fig. HPLC-ESI-MS/MS of bullatanocin from the alkaloid and acetogenin-rich fraction from *Annona crassiflora* fruit peel (AF-Ac) (*m*/*z* 661 [M+Na]^+^).**

**S12 Fig. HPLC-ESI-MS/MS of bullatacin/squamocin from the alkaloid and acetogenin-rich fraction from *Annona crassiflora* fruit peel (AF-Ac) (*m*/*z* 645 [M+Na]^+^).**

**S13 Fig. HPLC-ESI-MS/MS of annomontacin from the alkaloid and acetogenin-rich fraction from *Annona crassiflora* fruit peel (AF-Ac) (*m*/*z* 647 [M+Na]^+^).**

**S14 Fig. HPLC-ESI-MS/MS of desacetyluvaricin/isodesacetyluvaricin from the alkaloid and acetogenin-rich fraction from *Annona crassiflora* fruit peel (AF-Ac) (*m*/*z* 629 [M+Na]^+^).**

**S15 Fig. Cell viability of fibroblasts cells treated with the alkaloid and acetogenin-rich fractions of *Annona crassiflora* fruit peel (AF-Ac) or vehicle (control, cells treated with RPMI-1640 medium containing 0.05% DMSO). Results (mean ± SD, n = 3) expressed as the percentage of viable cells compared to the vehicle group. Significance levels are indicated by \**p* < 0.05 and \*\*\**p* < 0.001 when compared to control (one-way ANOVA and Dunnett as posttest).**

**The fibroblast cells (NIH/3T3) were grown in RPMI-1640 medium supplemented with fetal bovine serum 10% (FBS), 2 mM glutamine, 100 U/mL penicillin and 100 mg/mL streptomycin, at 37 °C and CO_2_ 5%. 3 x 10^4^ cells were plated in 96-well plates and treated with different concentrations of the AF-Ac fraction or vehicle and incubated for 24 h at 37 °C and 5% CO_2_. Then, 100 µL of 5 mg/mL (3-(4,5-dimethylthiazolyl-2)−2,5-diphenyltetrazolium bromide) solution was incubated with the supernatant at 37 °C for 2 h in 5% CO_2_. Next, dimethyl sulfoxide (DMSO) was added and the cell viability was analyzed by absorbance of the purple formazan from viable cells at 570 nm (Molecular Devices, Menlo Park, CA, USA).**

**S16 Fig. Western blot gels: six samples of each treatment group were evenly distributed in two gels resulting in 3 samples/gel/group. It was necessary to crop the membranes in order to incubate them with different antibodies since different proteins were analyzed in each western blot. Vinculin (124 kDa MW), β-actin (42 kDa MW) and PCNA (36 kDa MW) were analyzed in the same western blot by cutting the membrane in three (A). The top part was used to blot vinculin antibodies, the middle part was used to blot β-actin antibodies and the bottom part for PCNA antibodies. EGFR (180 kDa MW) and β-actin (42 kDa MW) were analyzed in the same western blot by cutting the membrane in two (B). The top part was used to blot EGFR antibodies and the bottom part for β-actin antibodies. The original blots show results of vinculin, β-actin, PCNA and EGFR expression of HepG2 cells treated with vehicle (control, showed in the three first lanes, included in the present study), crude ethanol extract from *A. crassiflora* fruit peel (EtOH, not included in the present study), n-butanol fraction from *A. crassiflora* fruit peel (BuOH, not included in the present study) and alkaloid and acetogenin-rich fraction from *A. crassiflora* fruit peel (AF-Ac, showed in the last three lanes, included in the present study). The corresponding MW (KD) markers are shown to the left of the Western blot image.**

